# A convenient single-cell newly synthesized transcriptome assay reveals *FLI1* downregulation during T-cell activation

**DOI:** 10.1101/2024.08.22.609222

**Authors:** Jun Lyu, Xiaoyan Xu, Chongyi Chen

## Abstract

Sequencing newly synthesized transcriptome alongside regular transcriptome in single cells enables the study of gene expression temporal dynamics during rapid chromatin and gene regulation processes. Existing assays for profiling single-cell newly synthesized transcriptome often require specialized technical expertise to achieve high cellular throughput, limiting their accessibility. Here, we developed NOTE-seq, a method for simultaneous profiling of regular and newly synthesized transcriptomes in single cells with high cellular throughput. NOTE-seq integrates 4-thiouridine labeling of newly synthesized RNA, thiol-alkylation-based chemical conversion, and a streamlined 10X Genomics workflow, making it accessible and convenient for biologists without extensive single-cell expertise. Using NOTE-seq, we investigated the temporal dynamics of gene expression during early-stage T-cell activation, identified transcription factors and regulons in Jurkat and naïve T cells, and uncovered the down-regulation of *FLI1* as a master transcription factor upon T-cell stimulation. Notably, topoisomerase inhibition led to the depletion of both topoisomerases and FLI1 in T cells through a proteasome-dependent mechanism driven by topoisomerase cleavage complexes, highlighting potential complications topoisomerase-targeting cancer chemotherapies could pose to the immune system.

## Introduction

Gene expression is a dynamic process involving the synthesis of new RNA transcripts and the degradation of old ones. Regular RNA sequencing methods cannot distinguish newly synthesized RNA from pre-existing RNA, thus unable to dissect transcription kinetics during biological processes involving rapid temporal changes in chromatin regulation and gene expression. Computational approaches, such as RNA velocity analysis, estimate RNA kinetics from existing RNA sequencing datasets by distinguishing unspliced and spliced mRNA molecules[1, 2]. However, these methods were based on certain assumptions that may lead to potential pitfalls[3]. Experimental approaches, which involve the metabolic labeling of newly synthesized RNA using nucleoside analogs followed by biochemical isolation or chemical conversion based on the incorporated nucleoside analogs, offer the capability to distinguish newly synthesized RNA from pre-existing RNA. This distinction enables accurate quantification of RNA synthesis and degradation rates[4]. Building on this strategy, several methods to detect and sequence newly synthesized RNA have been developed and applied to cultured cells in different settings, advancing our understanding of the regulation of transcription and RNA kinetics[5–9].

More recently, the strategy of metabolic labeling of newly synthesized RNA has been integrated into single-cell RNA sequencing[10], enabling the characterization of RNA kinetics with higher sensitivity, allowing for the detection of subtle temporal alterations in the transcriptome of single cells, and providing higher resolution in cell clustering and cell-fate trajectory analysis[11–18]. However, the applications of single-cell newly synthesized transcriptome sequencing to address scientific questions are not yet widespread, largely due to the technical limitations of existing methods. Plate-based assays, including scSLAM-seq[11], NACS-seq[12], and scEU-seq[13], have limited cellular throughput and potential batch-to-batch variations between individual plates. Among methods that offer high cellular throughput, sci-fate suffers from a low cell recovery rate of less than 5% due to the lengthy split-pool procedures[14], while scNT-seq, SLAM-Drop-seq, and Well-TEMP-seq all require specialized microfluidic systems or customized micro-well chips [15–18], limiting their accessibility to regular biologists without specialized single-cell technical expertise. On the other hand, the 10X Genomics platform is widely available in core facilities and often accessible to regular biology laboratories. The 10X Genomics workflow offers high cellular throughput, excellent performance, and convenience for single-cell transcriptome analysis with its streamlined protocol and automation, but lacks the capability of probing newly synthesized transcriptome in single cells.

In this study, we developed a new method, the **N**ewly synthesized transcriptome **o**n **10**X **E**xpression sequencing (NOTE-seq) assay, which integrates newly synthesized RNA detection into the 10X Genomics workflow for simultaneous profiling of both transcriptome and newly synthesized transcriptome in single cells. NOTE-seq retains the streamlined 10X Genomics workflow and data analysis pipeline, making it accessible and convenient for regular biologists without extensive experience in single-cell technology. Using NOTE-seq, we characterized the temporal dynamics of gene expression during early-stage T-cell activation, revealing patterns of transcription activities and gene regulation by transcription factors at different time points following T-cell stimulation. We identified *FLI1* as a master transcription factor with many downstream genes in its regulon, whose downregulation plays an important role in gene expression temporal dynamics upon T-cell stimulation. Additionally, FLI1 was depleted in T cells together with DNA topoisomerases upon topoisomerase inhibition. This protein depletion was induced by topoisomerase cleavage complexes via a proteasome-dependent mechanism, highlighting potential complications of the immune system that might be caused by certain types of DNA topoisomerase inhibitors widely used in cancer chemotherapy.

## Results

### NOTE-seq enables simultaneous profiling of transcriptome and newly synthesized transcriptome in single cells

The NOTE-seq workflow included metabolic labeling of the newly synthesized RNA, specific alkylation of the labeled nucleoside, and a modified protocol to profile single-cell transcriptome on the 10X Genomics platform (Fig. 1a). Specifically, after incubation with the thymidine analog 4-thiouridine (4sU) to label newly synthesized RNA, cells were formaldehyde crosslinked to immobilize RNA molecules in situ and iodoacetamide (IAA) treated to achieve thiol-specific alkylation of the incorporated 4sU, leading to the conversion from the thymidine analog to a cytosine analog, thus an effective T-to-C substitution in downstream sequencing analysis. Single-cell transcriptome profiling was then performed on the 10X Genomics platform following a modified workflow, including reverse transcription (RT) with cellular barcoding, reverse crosslink, and a second round of RT to enhance template-switching efficiency before cDNA amplification, followed by library preparation and sequencing. Generally speaking, sequencing reads containing T-to-C substitution represented newly synthesized RNA molecules, while those without T-to-C substitution represented pre-existing RNA. However, considering potential confounders arising from the relatively infrequent events of 4sU incorporation and sequencing errors, the exact fractions of newly synthesized RNA and pre-existing RNA were estimated based on a binomial mixture statistical model[19].

**Figure 1.**
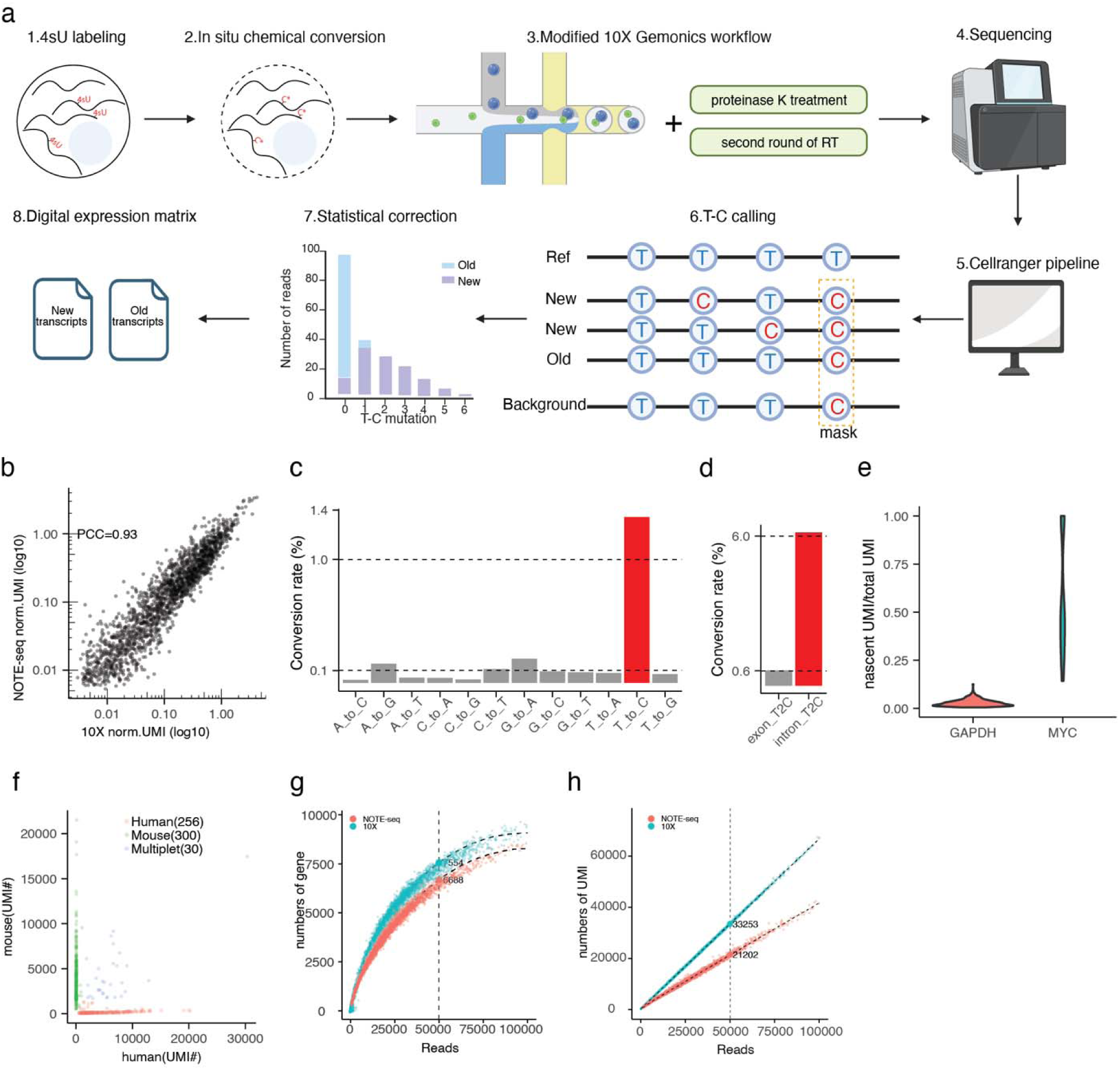
Workflow and performance of NOTE-seq. **a**. Workflow of NOTE-seq. **b**. Correlation of average normalized UMI counts of eHAP between NOTE-seq (n=3046 cells) and the standard 10X Genomics workflow (n=2774 cells). The Pearson’s correlation coefficient (PCC) was calculated based on 14771 genes. **c**. Overall base conversion rates derived from the NOTE-seq data, with the T-to-C conversion rate highlighted in red. **d**. T-to-C conversion rate in exonic reads and intronic reads. **e**. Fraction of newly synthesized RNA in a low-turnover-rate gene (GAPDH, n=674 cells) and a high-turnover-rate gene (MYC, n=128 cells). **f**. Transcript counts of individual cells mapped to the merged human-mouse genome, with cells colored based on their assigned categories of human, mouse, or multiplet. **g**. Comparison of detected gene numbers per cell between NOTE-seq and the standard 10X Genomics workflow. **h**. Comparison of detected UMI numbers per cell between NOTE-seq and the standard 10X Genomics workflow.

To characterize the performance of NOTE-seq, we carried out both NOTE-seq and regular single-cell transcriptome sequencing on the same 10X Genomics platform using the same human cell line eHAP1, and compared their parameters of single-cell transcriptome profiling. Comparable library size distributions (Fig. S1a) and similar fractions of mitochondrial and ribosomal genes (Fig. S1b) were observed, suggesting negligible RNA leakage that could skew RNA composition in the NOTE-seq workflow involving additional steps of 4sU labeling and subsequent thiol-alkylation-based chemical conversion. Moreover, single-cell transcriptomes characterized by both methods were highly correlated (Fig. 1b), suggesting additional steps in the modified NOTE-seq workflow did not significantly affect cell physiology, or alter gene expression patterns.

We further determined NOTE-seq’s specificity in labeling and identifying newly synthesized RNA. First, in the NOTE-seq data, the rate of T-to-C conversion was more than 10-fold higher than other types of conversion events (Fig. 1c), a strong indicator of successful 4sU incorporation and specific chemical conversion. Next, T-to-C events were mostly detected in intronic regions, with a 10-fold higher frequency than exonic T-to-C events (Fig. 1d), suggesting specific labeling of newly synthesized nascent RNA molecules instead of pre-existing RNA. Moreover, for the housekeeping gene *GAPDH* with a low transcript turnover rate, indeed only a small fraction of its RNA was newly synthesized, that was opposite to the *MYC* gene with a high transcript turnover rate and a dominant fraction of newly synthesized RNA (Fig. 1e), further confirming the specificity of NOTE-seq in detecting newly synthesized RNA. Finally, we analyzed single cells in different cell-cycle stages in the population. After confirming that NOTE-seq does not significantly alter the cell-cycle distribution (Fig. S2a), we focused on cell cycle markers in single cells within each stage. Consistent with the long mRNA half-life in human cells[20], we found that the detection of newly synthesized RNA often preceded the emergence of regular RNA during cell cycling (Fig. S2b), again demonstrating NOTE-seq’s capability and specificity to probe newly synthesized RNA in single cells.

As the cost of detecting newly synthesized transcriptome in addition to the regular transcriptome in single cells, NOTE-seq had some drawbacks compared to regular single-cell transcriptome sequencing on the 10X Genomics platform without distinguishing between newly synthesized and pre-existing RNA. First, cDNA length and the mapping ratio of sequencing reads were often compromised (Fig. S1c-d), likely due to the reduced processivity of the reverse transcriptase due to formaldehyde crosslinking and the chemical conversion of 4sU-incorporated RNA bases[15]. Consistently, NOTE-seq also showed a slight reduction in the detection sensitivity estimated by the number of genes and UMIs detected in single cells (Fig. 1g-h). In addition, NOTE-seq reported an elevated collision rate between single cells (Fig. 1f), yet still within the reasonable range, that was commonly observed in single-cell transcriptome assays involving formaldehyde crosslinking[21].

As a conclusion, we developed the NOTE-seq assay that enables simultaneous profiling of transcriptome and newly synthesized transcriptome in single cells. The slightly compromised gene detection sensitivity and elevated collision rate of the NOTE-seq assay compared to regular single-cell transcriptome analysis were largely tolerable in most scientific applications. Importantly, relying on the 10X Genomics workflow to offer high cellular throughput, NOTE-seq was a convenient and accessible method for regular biology labs lacking specialized expertise in single-cell technology. As a result, NOTE-seq was particularly suitable for biologists without extensive experience in single-cell assays and microfluidic automation, enabling them to conveniently examine single-cell transcriptome and newly synthesized transcriptome, which is crucial to understand the temporal dynamics of gene expression in single cells during processes involving rapid chromatin and gene regulation.

### NOTE-seq reveals gene expression temporal dynamics during T-cell activation

Using NOTE-seq, we studied T lymphocytes, a vital component of the immune system exerting central functions against infectious diseases and tumors. In particular, we investigated the rapid dynamics of gene expression during early-stage T-cell activation, a prerequisite for T-cell functions including T-cell development, proliferation, and differentiation[22]. In vivo, T cells can be activated via rapid stimulation of the canonical T-cell activation signaling pathway upon cognate antigen recognition. In vitro, the human Jurkat T cells provide a robust system to examine the temporal dynamics of gene expression during T-cell activation, after its stimulation by phorbol 12-myristate 13-acetate (PMA) and ionomycin (ION) to activate the downstream of the T-cell activation signaling pathway (Fig. 2a).

**Figure 2.**
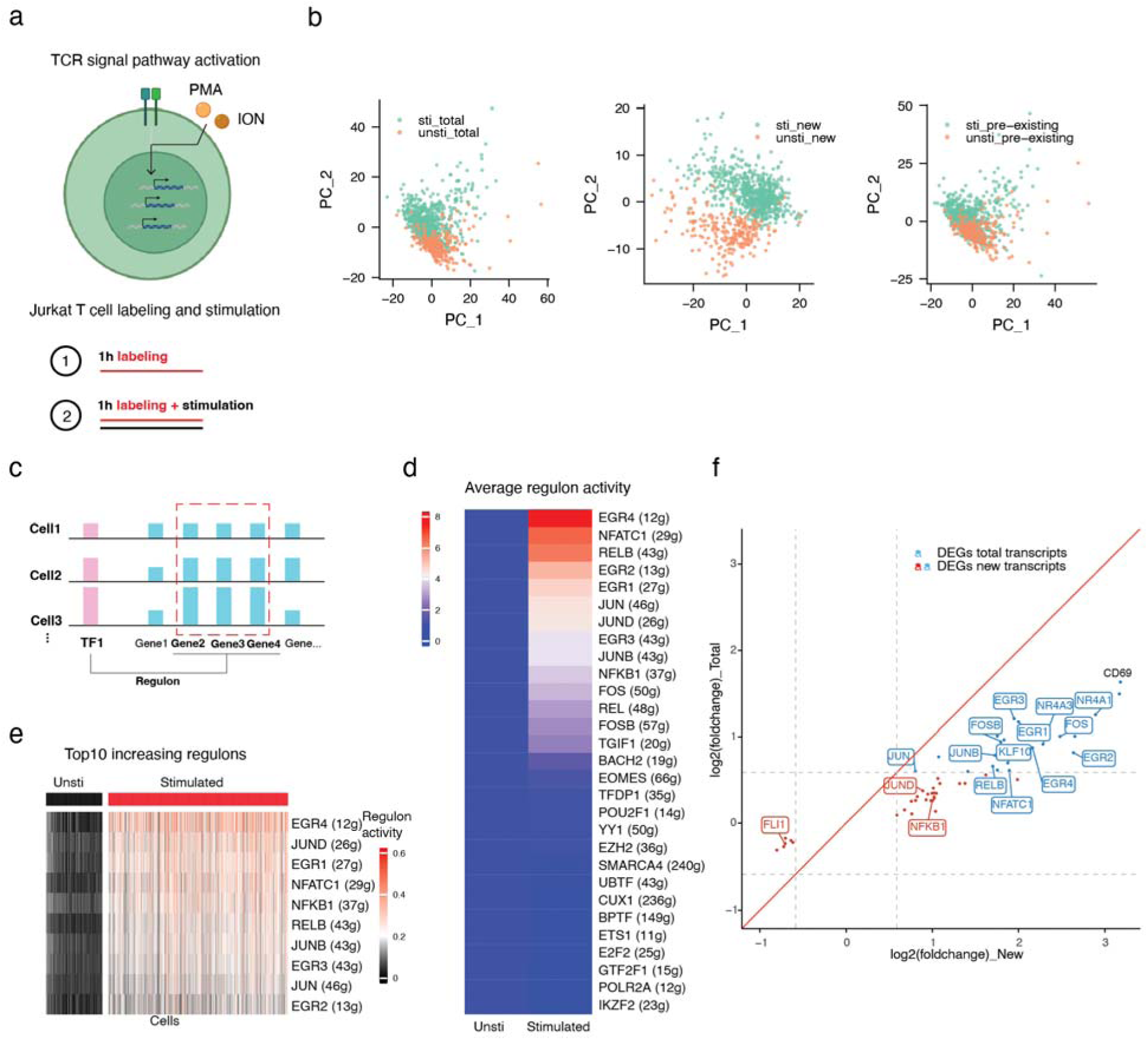
NOTE-seq reveals gene expression temporal dynamics during T-cell activation. **a**. Schematic of PMA/ION-mediated T-cell activation and 4sU labeling upon T-cell stimulation. **b**. Cell clustering analysis of stimulated (n=748 cells) and unstimulated (n=233 cells) Jurkat T cells by PCA, using total transcripts, newly synthesized transcripts (new), and pre-existing transcripts. **c**. Schematic of regulon analysis by correlating expression levels of transcriptional factors and their downstream genes. **d**. Comparison of regulon activities between unstimulated and stimulated Jurkat T cells. The stimulated regulon activity was normalized to the corresponding unstimulated regulon activity. **e**. Top 10 significantly activated regulons upon T-cell stimulation plotted for each single cell, with data derived from single-cell transcriptome matrix and scaled for each row. **f**. Differentially expressed genes identified using total transcripts or newly synthesized transcripts (blue) and only using newly synthesized transcripts (red) upon T-cell stimulation. Among the genes, transcription factors were highlighted with their names labeled in the boxes.

We characterized the temporal dynamics of gene expression during early-stage Jurkat T-cell activation using NOTE-seq (Fig. 2a). We found that the T-to-C conversion rate, the detection rate of newly synthesized RNA, the fraction of newly synthesized RNA per gene, and the sequencing depth per cell all showed a good similarity between conditions of stimulated T-cell activation and unstimulated negative control (Fig. S3a-d), suggesting the successful implementation of the NOTE-seq assay to investigate T-cell activation.

First, we performed cell clustering analysis. Principal component analysis (PCA) based on the single-cell transcriptome grouped individual cells into two partially overlapped clusters, indicating a lack of clear separation between stimulated and unstimulated cells (Fig. 2b, left panel). Next, we performed PCA based on the newly synthesized transcriptome revealed by NOTE-seq and successfully identified two clearly separated clusters, representing the stimulated and unstimulated T cells, respectively (Fig. 2b, middle panel). The better clustering performance could be attributed to the possibility that newly synthesized transcriptome in single cells, compared to regular transcriptome, more accurately reflects the difference between stimulated and unstimulated cells. We further reasoned that, considering the rapid gene expression dynamics upon T-cell activation and the relatively long RNA lifetime, the dominant fraction of the single-cell RNA could be pre-existing RNA transcribed before T-cell stimulation. Indeed, we found that PCA based on the pre-existing transcriptome generated two clusters that were mostly overlapped and indistinguishable (Fig. 2b, right panel). These results highlighted the importance of detecting the newly synthesized transcriptome by NOTE-seq, that was often missing in regular single-cell transcriptome assays, especially when studying dynamic processes involving rapid gene expression changes that could determine cell type and cell fate.

Next, we performed gene regulatory network analysis based on the temporal fluctuations of the transcriptome in single cells[23]. Specifically, after quantifying the transcription level of each gene that fluctuates in individual cells within the cell population, gene regulatory networks (a.k.a. regulons) were inferred by correlating the expression level of transcription factors (TFs) and the transcription level of their corresponding downstream targeted genes in single cells (Fig. 2c). Based on the NOTE-seq data, we identified multiple regulons corresponding to major T-cell activation signaling pathways in stimulated T cells (Fig. 2d-e), in agreement with the previous knowledge[22].

Finally, we investigated differentially expressed genes (DEGs) during T-cell activation (Fig. 2f). Not surprisingly, many canonical T-cell activation makers such as *CD69*[24] were upregulated at the RNA level. Moreover, we often noted a more significant increase in the level of newly synthesized RNA compared to the level of regular RNA (Fig. 2f), suggesting a superior DEG detection sensitivity by probing the newly synthesized RNA in NOTE-seq. Indeed, based on the level of newly synthesized RNA, we identified additional DEGs that otherwise would go unnoticed on the regular RNA level (Fig. 2f). Focusing on differentially expressed TFs during T-cell activation (Fig. 2f, Table S1), we found that most TFs with reported relevance in T-cell function were upregulated (Table S1), including families of *NR4A*, *EGR*, and *NF*κ*B*[25, 26]. Notably, we also identified a downregulated TF, *FLI1*, whose exact role during T-cell activation remains unclear.

### NOTE-seq reveals progressing transcriptome alterations and the TF turnover during naïve T-cell activation

The circumstances during T-cell activation in vivo may not be fully recapitulated by Jurkat T-cell activation in vitro. For example, naïve T cells are often quiescent in vivo, characterized by a small cell size, a low metabolism rate, and a low transcription activity[27], that is very different from the proliferating Jurkat T cells (Fig. 3a). Consequently, quiescence exit is an essential feature of naïve T-cell activation, together with other potential differences between naïve and Jurkat T cells, that could lead to a different gene expression program regulated by a different set of TFs.

**Figure 3.**
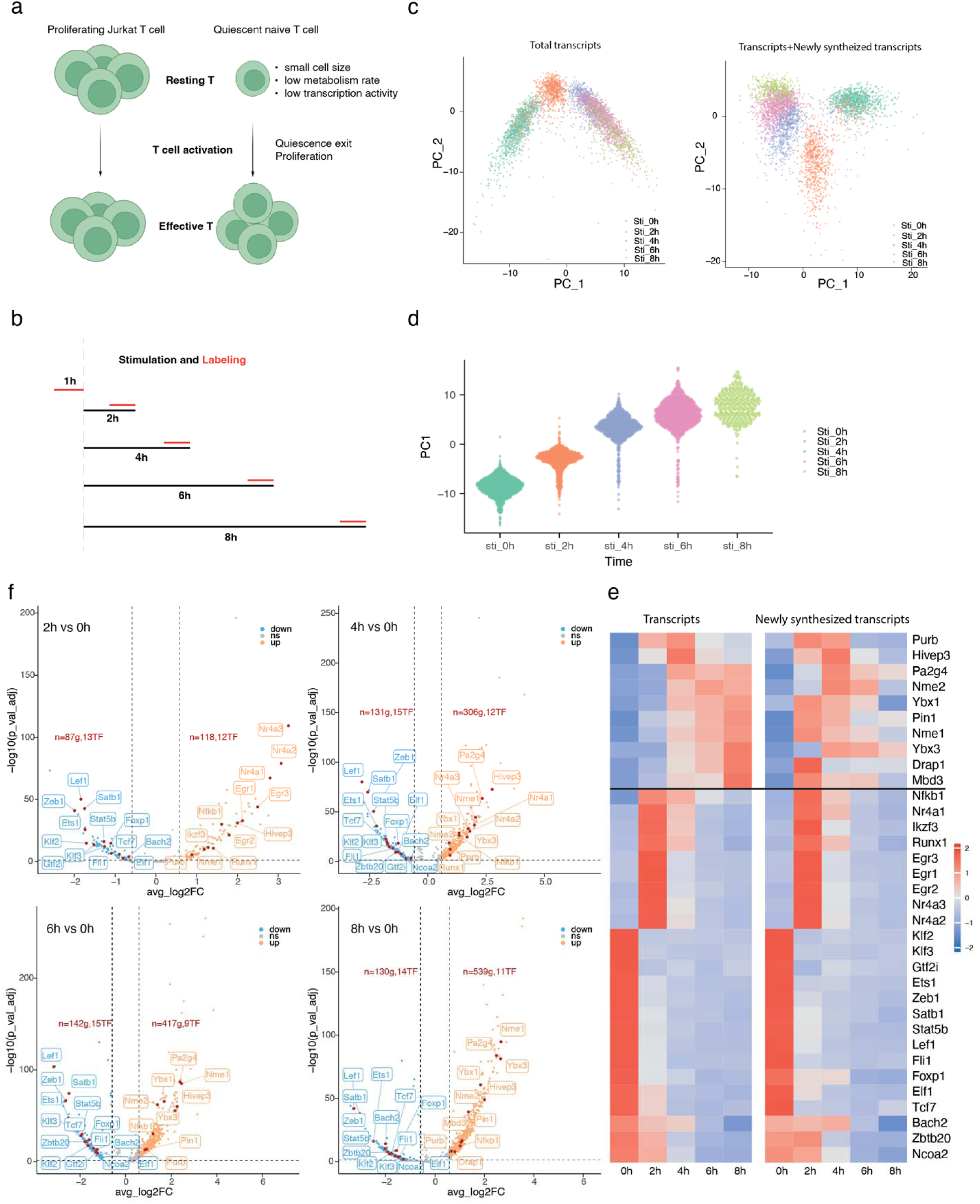
NOTE-seq reveals progressing transcriptome alterations and the TF turnover during naïve T-cell activation. **a**. Schematic of distinct processes during T-cell activation for Jurkat and naïve T cells. **b**. Strategy of naïve T-cell stimulation and 4sU labeling. Black line represents time for stimulation and the red line represents time for 4sU labeling. **c**. Cell clustering analysis of naïve T cells stimulated for the indicated time (1014 cells for 0 hour, 809 cells for 2 hours, 831 cells for 4 hours, 1016 cells for 6 hours, and 359 cells for 8 hours). PCA analyses and plots were based on total transcripts (left) or total and newly synthesized transcripts jointly (joint transcriptome, right). **d**. Principal components of the joint transcriptome at different time points after T-cell stimulation. **e**. Differentially expressed genes identified upon T-cell stimulation at different time points. Among the genes, transcriptional factors were highlighted with their names labeled in the boxes. **f**. Heatmap of the level of total transcripts and newly synthesized transcripts for each differentially expressed transcription factor, with the data matrix scaled for each row.

To address this concern, we characterized gene expression temporal dynamics during early-stage naïve T-cell activation ex vivo. Specifically, we isolated mouse naïve T cells for stimulation using agonistic antibodies and collected cells in a two-hour interval with one hour labeling of newly synthesized RNA prior to each harvest (Fig. 3b). The expected naïve T-cell activation was observed upon stimulation, as evidenced by the upregulation of multiple relevant makers (Fig. S4). The expected dose-dependence of T-to-C conversion frequency was also observed (Fig. S5a), suggesting successful 4sU labeling of newly synthesized RNA without triggering detrimental stresses or altering the gene expression program in naïve T cells (Fig. S5b-c).

First, we performed cell clustering analysis. Based on the single-cell transcriptome, we found that cells without stimulation, two-hour post-stimulation, and four-hour or longer post-stimulation were already well separated into distinct clusters (Fig. 3c), suggesting significant temporal alterations of the gene expression program upon naïve T-cell stimulation within the first four hours. Additional clustering based on both transcriptome and newly synthesized transcriptome offered a better resolution, and further separated cells at four, six, and eight hours post-stimulation into distinct clusters (Fig. 3c), suggesting minor yet distinguishable gene expression changes in the level of newly synthesized RNA after four hours post-stimulation. Furthermore, the first principal component of the single-cell joint transcriptomes demonstrated a temporal trend following the initial stimulation (Fig. 3d), again highlighting a dynamic program of gene expression progressing over time during naïve T-cell activation.

Next, we analyzed differentially expressed TFs over time during naïve T-cell activation, and identified different TFs at different time points (Fig. 3e, Table S2) after the initial stimulation. For many TFs, the changes in regular RNA level were often preceded by the changes in newly synthesized RNA level (Fig. 3f), which reflect the precise timing of the transcription response to T-cell stimulation. Some TFs showed similar trends between the levels of newly synthesized RNA and regular RNA at all time points (Fig. 3f), suggesting a high turnover rate for the transcripts of these TFs, a key feature of crucial gene regulators capable of fast modulation and acute response to the environmental stimuli.

Among the upregulated TFs identified during naïve T-cell activation, most have reported roles relevant to T-cell activation and function (Table S2), with many of them also coinciding with upregulated TFs observed during Jurkat T-cell activation, including families of *Nr4a*, *Egr*, and *Nf*κ*b*. The differences of a few TFs were likely due to differences in stimulation conditions, sampling time points, and T-cell origins. Interestingly, many downregulated TFs were identified specifically during the activation of naïve T cells but not Jurkat T cells (Fig. 3e). Many of them were involved in the maintenance of the quiescence state (Table S2), consistent with the additional essential step of quiescence exit during naïve T-cell activation.

Notably, *Fli1* was also downregulated during naïve T-cell activation (Fig. 3e), the same as our previous observation during Jurkat T-cell activation, suggesting *Fli1* as a newly identified TF with potential importance. Moreover, *Fli1* was downregulated rapidly at the start of T-cell activation, with a fast turnover rate since both its RNA level and newly synthesized RNA level rapidly respond to T-cell stimulation with the same trend (Fig. 3f), further indicating its role in regulating gene expression upon T-cell stimulation.

### *Fli1* is a master transcription factor downregulated during naïve T-cell activation

To further investigate the role of *Fli1* as a down-regulated TF during T-cell activation, we performed gene regulatory network analysis for the identification of master TFs that regulate a large number of downstream genes. Among the identified master TFs (Fig. 4a), the transient upregulation of *Nf*κ*b* is a hallmark of T-cell activation, with their upregulated regulon promoting cell proliferation (Fig. S6)[22, 28]. The *Klf3* and *Stat1* have reported roles in the maintenance of T cell naivety (Fig. S6)[29, 30], with their downregulated regulons expected for the quiescence exit during naïve T-cell activation.

**Figure 4.**
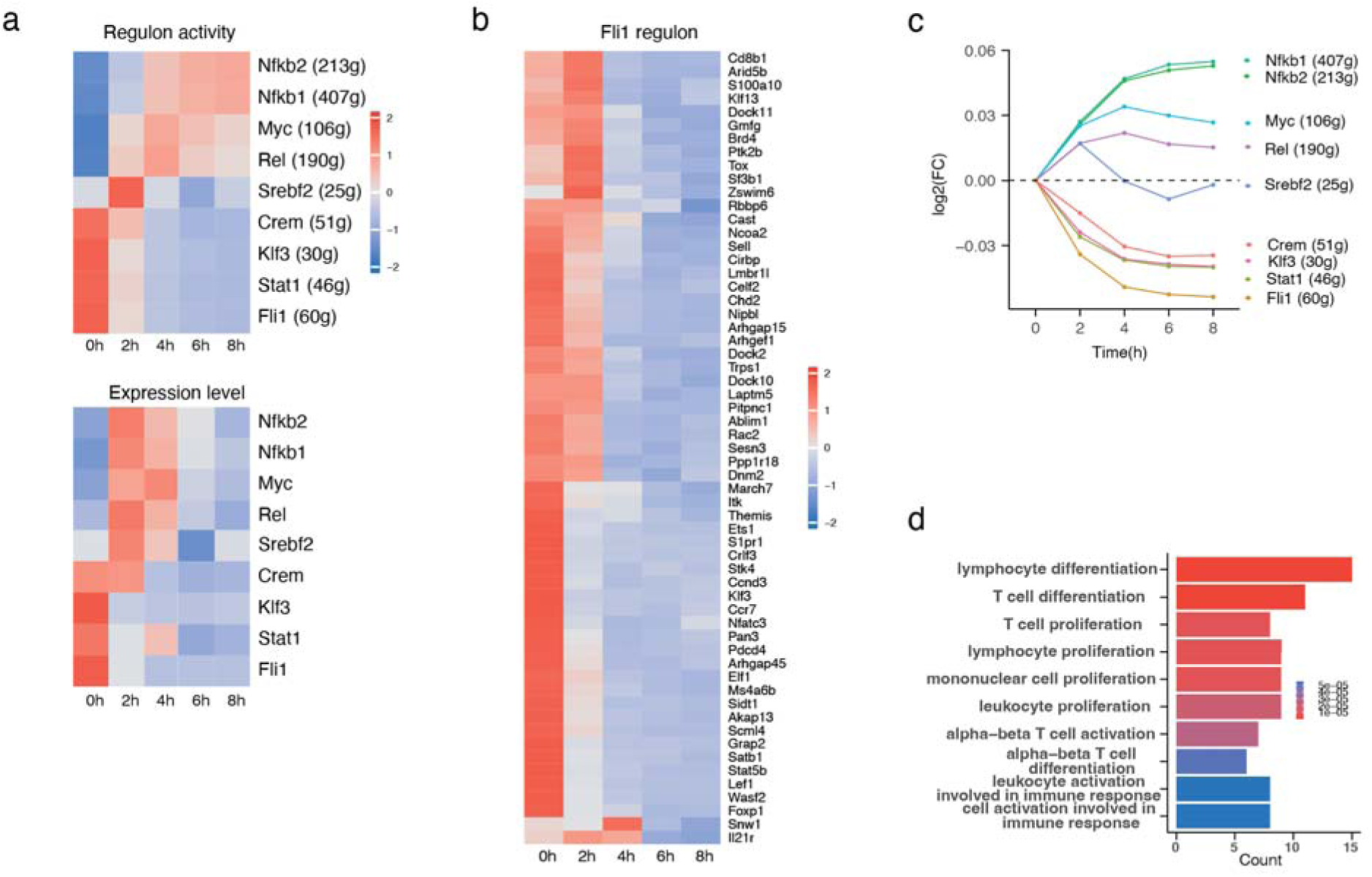
*Fli1* is a master transcription factor downregulated during naïve T-cell activation. **a**. Activity of regulons (top) and expression level of master TFs (bottom) at different time points during T-cell activation, with the data matrix scaled for each row. **b**. Transcription levels of downstream genes in the *Fli1* regulon at different time points during T-cell activation, with the data matrix scaled for each row. **c**. Fold changes of regulon activity at different time points during T-cell activation **d**. GO enrichment analysis of genes in the *Fli1* regulon.

*Fli1* was also identified as a master TF, regulating 59 downstream genes in the *Fli1* regulon (Fig. 4a). Most genes in the *Fli1* regulon were significantly downregulated over time (Fig. 4b), making *Fli1* regulon the most downregulated regulon among all the identified regulons during T-cell activation (Fig. 4c). Furthermore, gene ontology analysis showed that many genes in the *Fli1* regulon were categorized in T-cell related biological processes (Fig. 4d), suggesting a negative regulatory role of *Fli1* during T-cell activation, which echoes previous reports showing the inhibitory role of *Fli1* during T-cell activation and effector T-cell differentiation[31, 32].

### Topoisomerase inhibition induces proteasome-mediated FLI1 degradation in T cells

*FLI1* is a proto-oncogene[33] overexpressed in various types of human malignancies[34], most notably in Ewing sarcoma, which is characterized by *FLI1* translocation and the expression of the oncogenic fusion protein EWS/FLI1[34, 35]. To treat malignancies associated with FLI1 overexpression, small-molecule inhibitors of FLI1, including etoposide (ETO) and camptothecin (CPT) [34, 36], have been identified as effective treatments for Ewing sarcoma[36]. Both ETO and CPT are DNA topoisomerase inhibitors targeting human topoisomerase II (TOP2) and topoisomerase I (TOP1), respectively, leading to the catalytic inhibition of topoisomerase and the enrichment of topoisomerase cleavage complex (TOPcc).

We hypothesized that ETO and CPT may act as FLI1 inhibitors not only in specific malignancies but also generally in normal cells including T cells. Indeed, we found that both drugs induced a dose-dependent reduction of the FLI1 level in both naïve and Jurkat T cells (Fig. 5a-b). Such FLI1 depletion relied on the formation of TOPcc, since the same depletion was not observed upon ICRF-193 treatment that inhibits TOP2 catalytic activity without the formation of TOP2cc (Fig. 5b).

**Figure 5.**
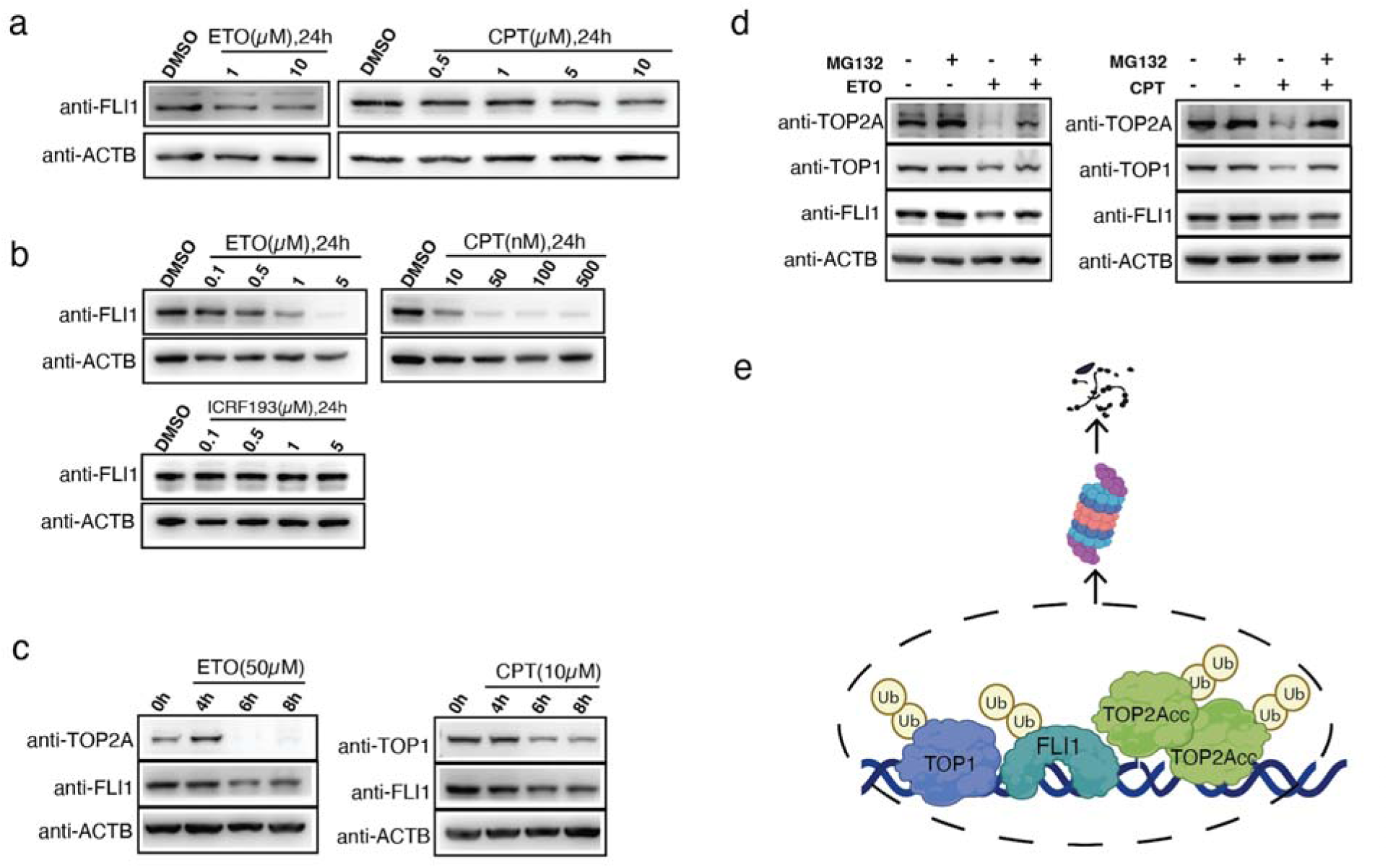
Topoisomerase inhibition induces proteasome-mediated FLI1 degradation in T cells. **a**. FLI1 depletion in naïve T cells after topoisomerase inhibition. **b**. FLI1 depletion in Jurkat T cells after topoisomerase inhibition and TOPcc formation. **c**. Co-depletion of FLI1 and topoisomerase upon TOPcc formation. **d**. Proteasome-depenent degradation of FLI1, TOP1, and TOP2A upon TOPcc formation. **e**. Model of TOPcc-induced proteasome-mediated degradation of the potential topoisome complex including FLI1, TOP1, and TOP2A.

We further studied the mechanism of TOPcc-dependent FLI1 depletion. We ruled out TOPcc-mediated transcription inhibition, as no FLI1 reduction was observed in the RNA level (Fig. S7a-b). It could not be attributed to secondary complications that TOPcc may cause during DNA replication and cell division, as FLI1 depletion was also observed within much shorter time scales (Fig. 5c). Interestingly, concomitant with the depletion of FLI1 upon ETO treatment and TOP2cc formation, we also observed the depletion of TOP2A and TOP1 (Fig. 5c-d). Similarly, upon the depletion of FLI1 during CPT treatment and TOP1cc formation, there was depletion of both TOP1 and TOP2A (Fig. 5c-d). Given that TOPcc formation triggers its resolution via proteasome degradation[37], and TOP2A and TOP1 were reported to be in the same topoisome complex[38], we hypothesized that the TOPcc-dependent depletion of FLI1, TOP1, and TOP2A may occur in the same protein complex in a proteasome-dependent manner. Indeed, we found that TOPcc-induced degradation of both topoisomerases and FLI1 was rescued by proteasome inhibition (Fig. 5d, Fig. S8).

Finally, we concluded that FLI1 depletion in T cells upon topoisomerase inhibition was induced by TOPcc via proteasome degradation of the potential FLI1-TOP1-TOP2A topoisome complex (Fig. 5e). Considering that both ETO and CPT derivatives are frontline cancer chemotherapeutic drugs targeting DNA topoisomerases[39, 40], our discovery indicated potential complications such chemotherapies may pose to the immune system beyond their main effects in tumor cells.

## Discussion

Single-cell newly synthesized RNA sequencing, in addition to regular single-cell RNA sequencing, allows simultaneous profiling of newly synthesized transcriptome and regular transcriptome in single cells, enabling the investigation of gene expression temporal dynamics during rapid chromatin and gene regulation. We developed the NOTE-seq assay that can be integrated into the streamlined 10X Genomics workflow, offering simultaneous quantification of both newly synthesized and pre-existing transcriptomes in single cells with high cellular throughput. Importantly, NOTE-seq is easily accessible to many biology laboratories, and can be readily performed by regular biologists without specialized expertise and extensive experience in single-cell technology.

Using NOTE-seq to characterize gene expression temporal dynamics upon T-cell stimulation, we identified TFs and their downstream genes forming regulons, and uncovered the downregulation of a new master TF *FLI1* during T-cell activation. However, potential applications of NOTE-seq extend far beyond the processes studied in this work. Generally speaking, NOTE-seq enables the investigation of any dynamic process involving rapid changes in chromatin and gene regulation over time at the single-cell level, as newly synthesized RNA accurately reflects the precise timing of transcription kinetics, that are often delayed or hidden in regular RNA analysis. Additionally, NOTE-seq quantifies the level of both regular RNA and newly synthesized RNA in single cells with high accuracy, that is crucial to estimate important parameters including RNA synthesis and degradation rates.

While the manuscript was under preparation, a conceptually similar method, NR-scRNA-seq, was reported, demonstrating a chemical-conversion-compatible 10X Genomics workflow to probe newly synthesized RNA in single cells[41]. However, unlike in the NOTE-seq assay that RNA molecules were anchored within cells via formaldehyde crosslink and fixation, this method directly probed RNA in methanol-fixed yeast cells, which may suffer from the RNA leakage issue. While yeast cells have cell walls that may reduce the outflow of RNA molecules thus alleviating this concern, the potential caveat of RNA leakage could significantly compromise its performance and RNA detection sensitivity in single mammalian cells without cell walls. A side-to-side comparison will be needed for further evaluation of these two methods.

It is intriguing that DNA topoisomerase inhibitors ETO and CPT, which lead to the formation of TOP2cc and TOP1cc respectively, reduce the levels of both topoisomerases and FLI1 in T cells. One possible scenario is the formation of a topoisome complex consisting of TOP1, TOP2A, FLI1, and possibly also involving other players such as MYC and p53 as reported[38, 42], and this complex could be co-degraded by the proteasome pathway that is activated upon TOPcc formation. Further studies will be needed to elucidate the molecular mechanism of FLI1 depletion upon topoisomerase inhibition.

In addition to its downregulation as a master transcription factor during T-cell activation, *FLI1* is also implicated in regulating T-cell development[34] and the response of effector T cells[32], underscoring a potentially broad impact of *FLI1* downregulation in the immune system upon topoisomerase inhibition. Indeed, various studies across different contexts have suggested important roles of DNA topoisomerases in immune diseases[43, 44], and that topoisomerase inhibition could modulate the immune response during inflammation[45, 46]. Further research is essential to deepen our understanding of the roles of FLI1 and the effects of cancer chemotherapeutic topoisomerase inhibition in T-cell function and the overall immune response, including exploring different T-cell subsets, various stages of T-cell activation, and other components of the immune system.

## Materials and Methods

### Cell culture

The eHAP human cells (Horizon discovery Ltd, Cat # C669) were cultured in IMDM medium (Gibco) containing 10% FBS and 1% PS, and supplemented with 1 mM 4sU one hour for RNA labeling before harvest. Jurkat T cells (E6.1 clone) were cultured in RPMI1640 medium (Gibco) containing 10% FBS and 1% PS. For RNA labeling and T-cell stimulation, one million Jurkat T cells were resuspended in RPMI1640 containing 250 µM 4sU, 81 nM phorbol 12-myristate 13-acetate (PMA) and 1.34 µM ionomycin (eBioscience™ Cell Stimulation Cocktail) for one-hour incubation. Naïve T cells were purified from mouse splenocytes and lymphocytes of C57BL/6J WT mice using EasyStem^TM^ Mouse Pan-Naïve T cell isolation kit (STEMCELL), and cultured in RPMI1640 medium containing L-glutamine, 50 µM 2-mercaptoethanol, 10% FBS, and 1% PS. For RNA labeling and T-cell stimulation, 0.5-1 million cells were placed in a 24-well plate coated with 5 ng/µl plate-coated anti-CD3 (Biolegend, clone 2C11) and anti-CD28 (Biolegend, clone 37.51) for incubation for the time indicated, followed by 500 µM 4sU addition one hour before harvest.

### Workflow of the NOTE-seq assay

One million 4sU-labeled cells were harvested, resuspended in 500 µl ice-cold PBS, fixed by adding 500 µl 8% formaldehyde, and incubated on ice for 15 minutes. After washing with PSD (0.5% Superase•In and 10 mM DTT in PBS), cells were resuspended with PSD containing 0.2% Triton X-100, incubated on ice for 5 minutes, washed with PS (0.5% Superase•In in PBS), and resuspended in 100 µl PS. Then, 40 µl fresh-made iodoacetamide (IAA), 40 µl 500 mM Sodium phosphate buffer (pH 8.0), and 20 µl nuclease-free water were sequentially added to the 100 µl cell suspension, before mixing with 200 µl DMSO. After 15-minute incubation at 50°C on a mixer, cells were put on ice with immediate addition of 8 µl 1 M DTT to quench the chemical conversion. Cells were then washed with 5 ml ice-cold PBS containing 0.2% Triton X-100 and resuspended in 200 µl PS. Cell suspension was loaded on the 10X Genomics platform, following the workflow according to the Chromium Next GEM Single Cell 3 Kit v3.1 user guide (Rev E) with the following modifications: (1) After the GEM break (step 2.1a), the aqueous phase was transferred to a new PCR tube, with the addition of 2 µl 20 mg/ml protease K (Invitrogen) and one-hour incubation at 53°C on a mixer. (2) After Dynabeads Cleanup (step2.1n), beads were resuspended with 10.5 µl MixtureA consisting of 50 mM Tris-HCl (pH 8.0), 60 mM NaCl, 5 mM MgCl_2_, and 1 mM dNTP. 10 µl elution was mixed with 10 µl MixtureB consisting of 2 mM GTP, 10% PEG8000, 16 mM DTT, 2U/µl Superase•In, 4 µM TSO, and 200 U Maxima H-minus, and incubated at 53°C for 45 minutes (second round of reverse transcription). (3) cDNA was amplified by mixing 50 µl Amp Mix, 15 µl cDNA primers, and 15 µl nuclease-free water, following the PCR program and cycles in the user guide (step2.2d). (4) The PCR products were purified twice, following step2.3. The downstream library preparation and sequencing were performed the same as the user guide.

### The barnyard experiment

Human Jurkart T cells and NIH3T3 cells were mixed at a 1:1 ratio, followed by the NOTE-seq workflow without T-to-C conversion. Reads were aligned to GRCh38_and_mm10-2020-A reference by cellranger7.1 with cellular barcodes assigned to human, mouse, or multiplets. Briefly, cellular barcodes are initially classified as human or mouse by which genome has more total UMI counts for that barcode. Barcodes with total UMI counts that exceed the 10th percentile of the distributions for both human and mouse are called observed multiplets. Because cellranger only detects the observable multiplets (human, mouse), it computes an inferred multiplet rate by estimating the total number of multiplets, including (human, human) and (mouse, mouse). This is done by estimating via maximum likelihood the total number of multiplet GEMs from the observed multiplets and the inferred ratio of human to mouse cells.

### scRNA-seq data analysis

Raw sequencing data was subjected to fastq calling, reads alignment, and quantification using cellranger7.1 with default parameters and standard pipeline. The gene expression matrix from the filtered_feature_bc_matrix was used to create a Seurat object using Seurat v4.3.0 with default parameters. Cellular barcodes (CBs) were retrieved from the Seurat object, and were used to subset the BAM file. Subsequently, confidently unique-aligned reads representing UMIs (corresponding to BAM tag: xf:i:25) were extracted from reads associated with the selected CBs. A modified pipeline of scNT-seq was adopted to quantify metabolically labeled and unlabeled transcripts. Briefly, sites of T-to-C substitution with the base Phred Quality Score > 27 (A to G substitution in the reverse complementary strand) in each transcript were extracted and then filtered by background control without 4sU labeling and chemical conversion. The resultant transcripts containing T-to-C substitution were counted as newly synthesized transcripts and then corrected by a statistical model described below.

### Estimation of the fraction of newly synthesized transcripts

The metabolic labeling of newly synthesized transcripts was not 100% complete due to insufficient 4sU substitution, resulting in underestimation of newly synthesized transcripts. To remedy such quantification bias, a binomial mixture model was adopted to approximate the real distribution of T-to-C substitution and the real mutation rate. Briefly, the likelihood function for each gene transcript i follows:

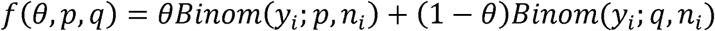

where θ is the fraction of newly synthesized transcripts in each experiment, p and q are the probabilities of a T-to-C substitution at each nucleotide for newly synthesized and pre-existing transcripts, respectively, and *n_i_* is the number of uridine nucleotides observed in the transcript i. As the p and q are global parameters (not gene-specific) for all transcripts, for each sample, 10,000 UMIs were randomly sampled to estimate the global p and q. The model was fit by maximizing the likelihood function using the Nelder–Mead algorithm. The optimization was repeated 100 times with random initialization values for θ, p, and q in the range [0,1]. The p and q were kept if θ is in the range [0,1]. The p and q corresponding to the maximal function value were used for calculating θ of individual genes with >= 100 UMIs, using the Brent algorithm with the constraint θ∈[10^−8^,1]. The labeled transcript levels of these genes (N_gene_) were re-estimated as follows:

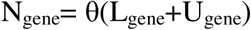

where θ is the fraction of newly transcribed transcripts for a gene, L_gene_ is the number of labeled transcripts of a gene and U_gene_ is the number of unlabeled transcripts of a gene.

Since the detection rates of labeled transcript (α) are highly similar in one cell, the average α was calculated for all genes in one cell as follows:

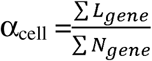

where L_gene_ is the number of observed labeled transcripts of a gene with estimated N_gene_ in that cell.

Then, the new transcript level for each gene in each cell can be re-calculated as follows:

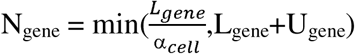

The statistic correction pipeline was adapted from Well-TEMP-seq[16].

### Cell cycle analysis

The normalized Seurat objects containing single-cell expression matrices were subjected to CellCycleScoring() function of Seurat (v4.3.0), where individual cells were scored, based on updated gene sets of cell cycle markers (cc.genes.updated.2019), and then classified into distinct cell cycle phases. The average expression levels of canonical cell cycle markers were visualized using heatmap[47].

### SLAM-seq

One million naïve T cells were incubated with varying concentrations of 4sU and stimulated by 5 ng/µl plate-coated anti-CD3 and anti-CD28 for 2 hours before being harvested for RNA extraction following the manual of RNeasy Mini Kit (Qiagen). One million Jurkat T cells were incubated with 50 nM CPT and 1 µM ETO for 24 hours with 1 hour 500 µM 4sU labeling before harvest. Next, RNA was subject to SLAM-seq protocol with minor modifications. Briefly, 2 µg RNA were combined with 10 mM iodoacetamide (Sigma-Aldrich, St. Louis, MO, USA), 50 mM sodium phosphate buffer (pH 8.0) and 50% DMSO, followed by 15 min of incubation at 50°C. The reaction was stopped by adding 20 mM DTT, followed by RNA purification using 1.8X RNA beads (Aline Bioscience) and RNA elution with 11 µl nuclease-free water. Then, 500 ng RNA was subject to library construction using Quant-seq 3′ mRNA-Seq Library Prep Kit (Lexogen). Data were first trimmed using cutadpt (v1.15) and then analyzed using SLAM-DUNK (v0.4.3)[48] to map, filter and quantify reads containing T-to-C mutation.

### PCA analysis, DEG identification and GO enrichment

Seurat objects were generated with newly synthesized, pre-existing, and total transcript matrices. Functions in the Seurat package including RunPCA and FindMarkers were used to perform principal component analysis (PCA) and identify differentially expressed genes (DEGs) using Wilcoxon test, respectively. Genes with absolute fold changes in expression > 1.5 and adjusted p-value < 0.05 were selected as DEGs. The list of transcription factors to start with was according to previous studies[49, 50]. Gene Oncology analysis and KEGG pathway analysis were performed by clusterProfiler[51].

### Gene regulatory network analysis

Regulon activity analysis was performed using single-cell regulatory network inference and clustering (SCENIC). Briefly, the newly synthesized transcript matrix and total transcript matrix were filtered with default parameters of geneFiltering, and the correlation matrix was computed based on gene expression level by runCorrelation. The correlation matrix of TFs and their target gene were computed and weighed by GENIE3, and then correlation modules were validated (with RcisTarget), selected (with weight > 0.005), and scored using runSCENIC. The resultant modules (regulons) with highly confident annotation for motifs of target genes were used for downstream analysis.

### Flow cytometry

Flow cytometry was performed in Beckman CytoFlex LX flow Cytometer, and data were analyzed with Flowjo v10 software (TreeStar). The following antibodies, conjugated with designated fluorochromes were used: anti-mouse CD69 (H1.2F3), CD62L (MEL-14), CD4 (RM4-5), CD8 (53-5.8), CD90.2 (53-2.1), CD44 (1M7), anti-phospho AKT (S473), anti-phospho-S6 (Ser235/236). Viakrome 808 Fixable Viability dye (Beckman Coulter) was used for viability staining for 20 min at room temperature. For cell surface staining, cells were stained with different antibody cocktails in a staining buffer (PBS with 5% FCS, 2 mM EDTA, 1ug/ml Fc receptor blocking antibody) for 30 min on ice, followed by viability staining and fixation with PFA. For intracellular staining, cells were fixed and permeabilized with Cytofix/CytopermTM solution (BD), followed by staining against intracellular antigens. For Phosflow staining, cells were fixed with BD Phosflow Fix Buffer for 10 min and permeabilized in BD Phosflow Perm buffer for 30 min on ice. Then cells were stained with indicated antibodies in staining buffers for 1 hours on ice.

### Western blot

To probe FLI1 depletion, naïve T cells and Jurkat T cells were treated with indicated concentrations of CPT and ETO for the indicated time. To probe FLI1 rescue, Jurkat T cells were treated with 20 µM MG132 and 50 µM ETO or 10 µM CPT for 6 hours. Then, cells were lysed in RIPA buffer supplemented with protease inhibitors for protein extraction, followed by 95°C protein denaturing in 1x SDS loading buffer. The equal volume of protein samples was loaded to and separated by 10% SDS-PAGE and then transferred onto an NC membrane (Millipore). The membrane was blocked in 5% non-fat milk dissolved in Tris-buffered saline with Tween-20 (TBST) for 1 hour and then washed with TBST three times. The membrane was then incubated with rabbit anti-mouse primary antibodies against TOP1(Abcam, ab109374), TOP2A (Abcam, ab12318), FLI1 (Abcam, ab133485) or beta-actin (Abcam, ab272085) overnight at 4°C, and with horseradish peroxidase-labeled secondary antibody (Abcam, catalog no. ab97080) at room temperature for 1 hour. The protein signals were detected by ECL Kit (Thermo Scientific, 34580) on Cytiva800.

### Sequencing

Libraries were sequenced on an Illumina NextSeq 2000 instrument. For NOTE-seq libraries, sequencing parameters were determined following the Chromium Next GEM Single Cell 3 Kit v3.1 user guide. For SLAM-seq libraries, 100 cycles-single-end sequencing was performed for read1.

## Data availability

All count tables generated by NOTE-seq pipeline are available at https://github.com/lyuj2022/NOTE-seq. The raw sequencing data were deposited at the National Center for Biotechnology Information Gene Expression Omnibus (GEO), with the accession number GSE270292.

## Code availability

The main scripts used to analyze data and generate main figures and conclusions are available at https://github.com/lyuj2022/NOTE-seq.

## Acknowledgements

We thank Madeline Wong and Steve Shema from the CCR Genomics Core for sequencing; Charlie Seibert from Single Cell Analysis Facility for 10X Genomics training; and other members of the Chen lab for discussions. The Jurkat T cell line is a gift from Dr. Peter D. Aplan. The mouse primary T cells are provided by Dr. Rachel Caspi. This study used the Biowulf Linux cluster at the National Institutes of Health. This work was supported by the Intramural Research Program of the National Institutes of Health, National Cancer Institute.

## Author contributions

J.L. and C.C. conceived the project and wrote the manuscript. J.L. developed the assay protocol, modified the data analysis pipeline, performed most of the experiments, analyzed most of the data, and prepared the figures. X.X. performed some experiments, analyzed some data, and provided helpful suggestions. C.C. supervised this project.

