## Supplemental Figure 1-8 for "A convenient single-cell newly synthesized transcriptome assay reveals *FLI1* downregulation during T-cell activation"

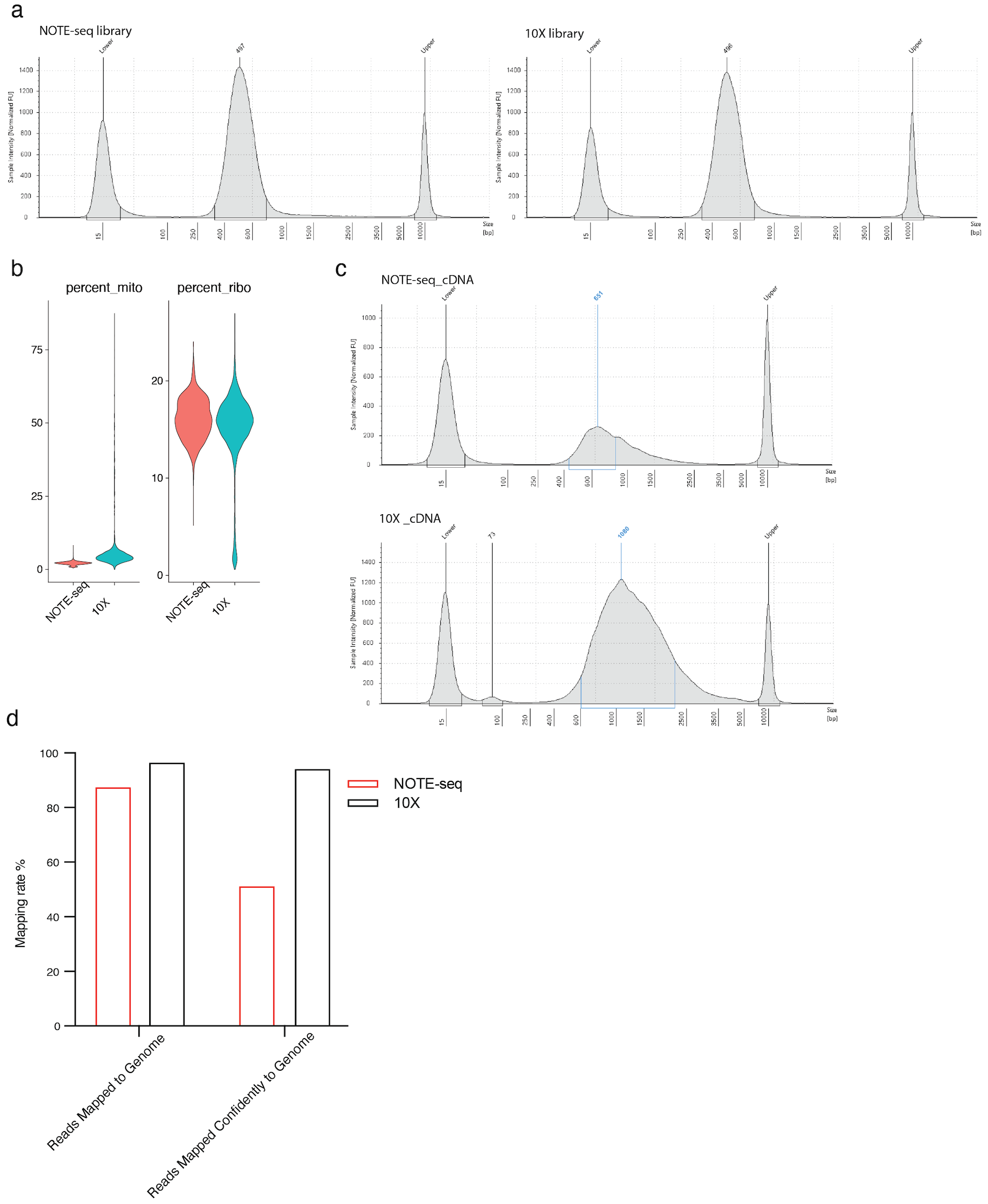

Figure S1. Comparison between NOTE-seq and the standard 10X Genomics workflow. **a**. Library length distribution of NOTE-seq and the standard 10X Genomics workflow. **b**. UMI percentages of mitochondrial genes and ribosomal genes in the data of NOTE-seq (n=3046 cells ) and the standard 10X Genomics workflow (n=2774 cells). **c**. cDNA length distribution of NOTE-seq and the standard 10X Genomics workflow. **d**. Reads mapping rate of NOTE-seq and the standard 10X Genomics workflow. Read mapped confidently to the genome denotes the fraction of reads mapped uniquely to the genome.

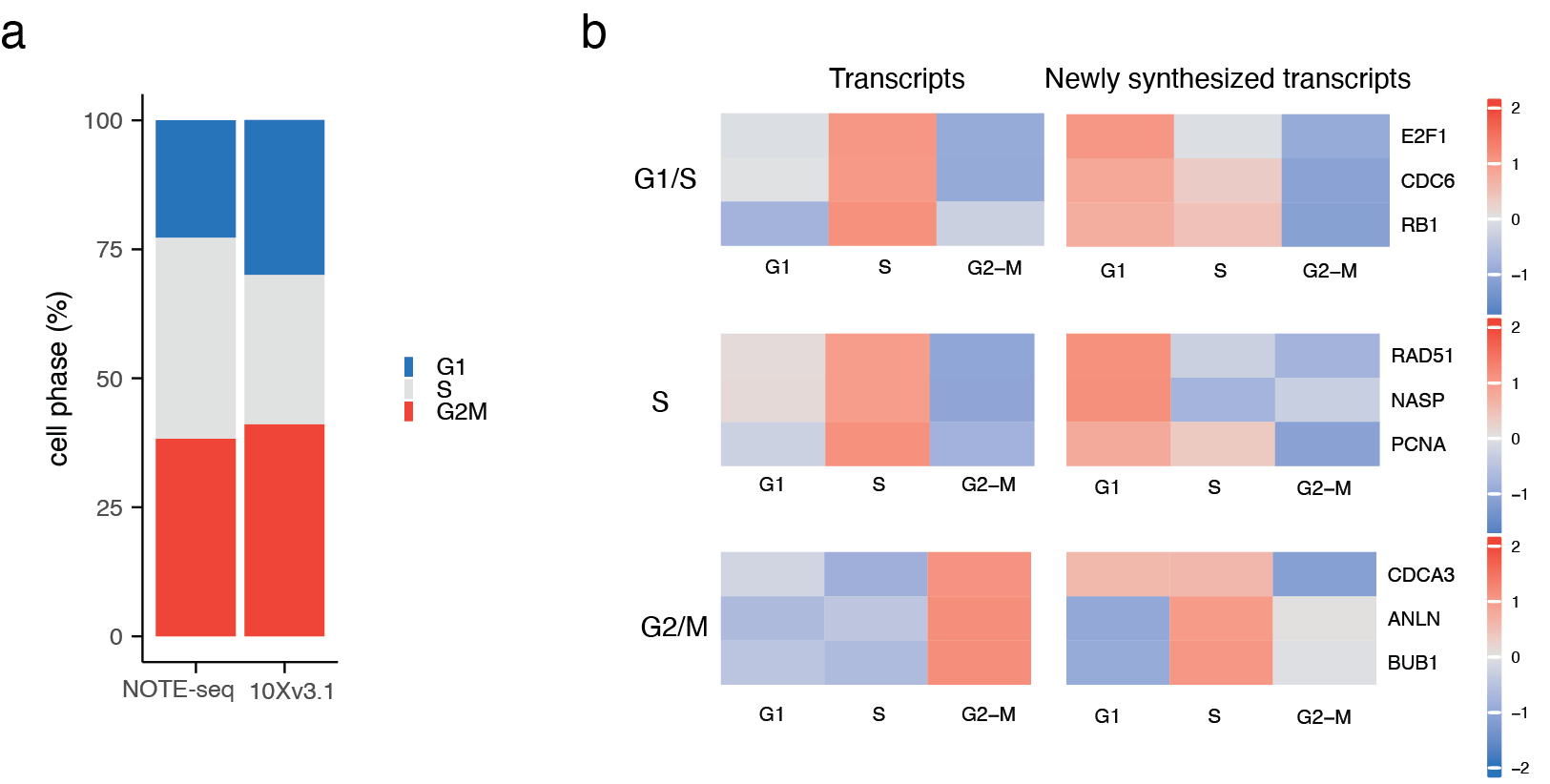

Figure S2. Cell cycle analysis using datasets from NOTE-seq (n=1378 cells) and the standard 10X Genomics workflow (n=1460 cells). **a**. Factions of cells in different cell-cycle stages. **b**. Transcription levels of cell-cycle marker genes at different cell-cycle stages, plotted by total transcripts and newly synthesized transcripts. The data matrix was scaled for each row.

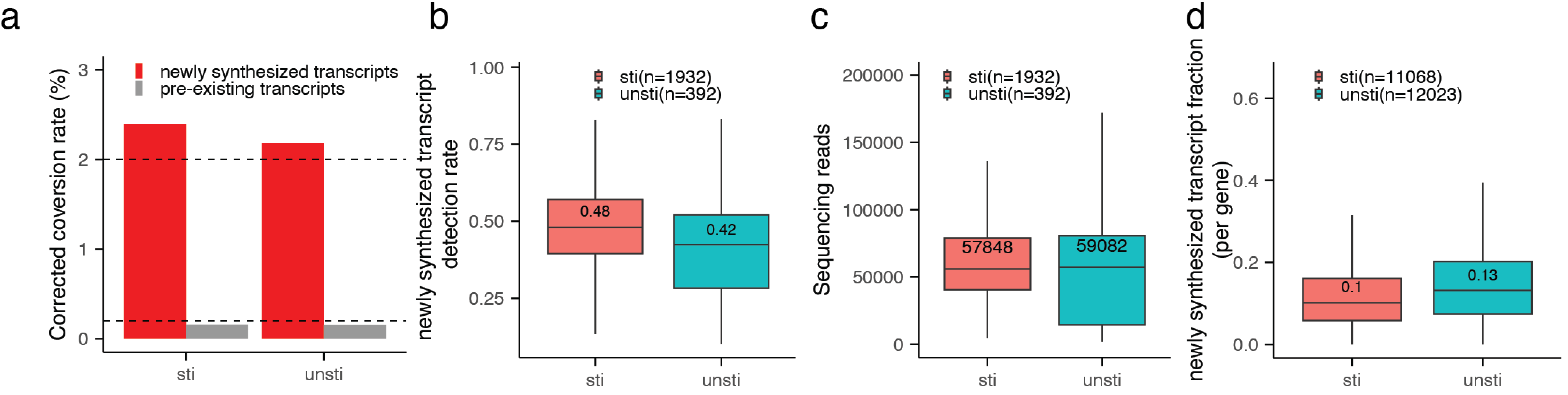

Figure S3. Comparison between unstimulated and stimulated Jurkat T cells. **a**. T-to-C conversion rates after statistical model correction. **b**. Distribution of newly synthesized RNA detection rates in each cell. **c**. Distribution of sequencing depth of each cell. **d**. Distribution of the fraction of newly synthesized RNA for each gene.

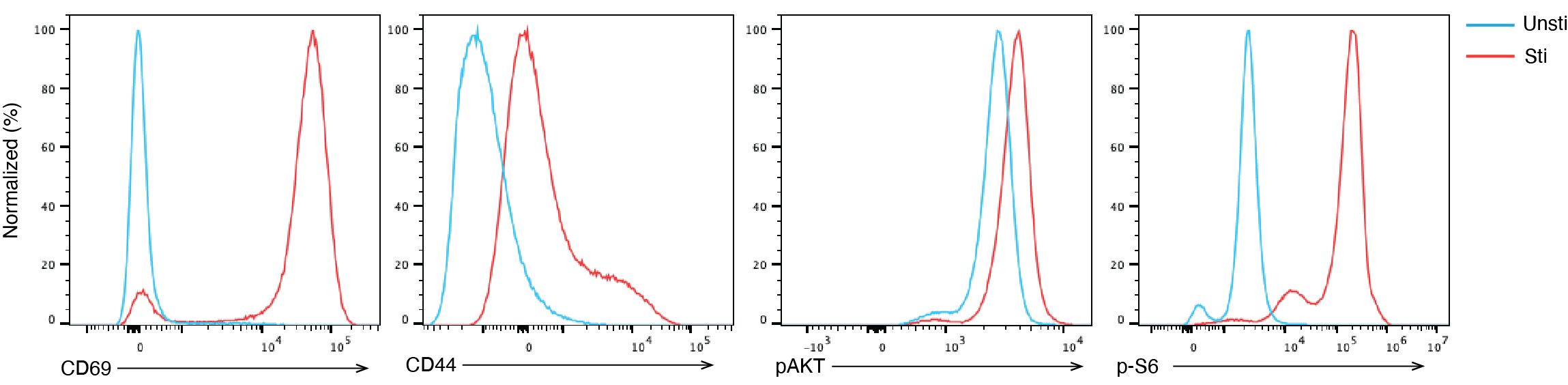

Figure S4. Naïve T-cell stimulation and activation by agnostic antibodies. Flow cytometry data showing the upregulation of markers representing T-cell activation (CD69 and CD44) and quiescence exit (pAKT and p-S6), after 8-hour stimulation by 5µg/µl plate-bound anti-CD3/CD28.

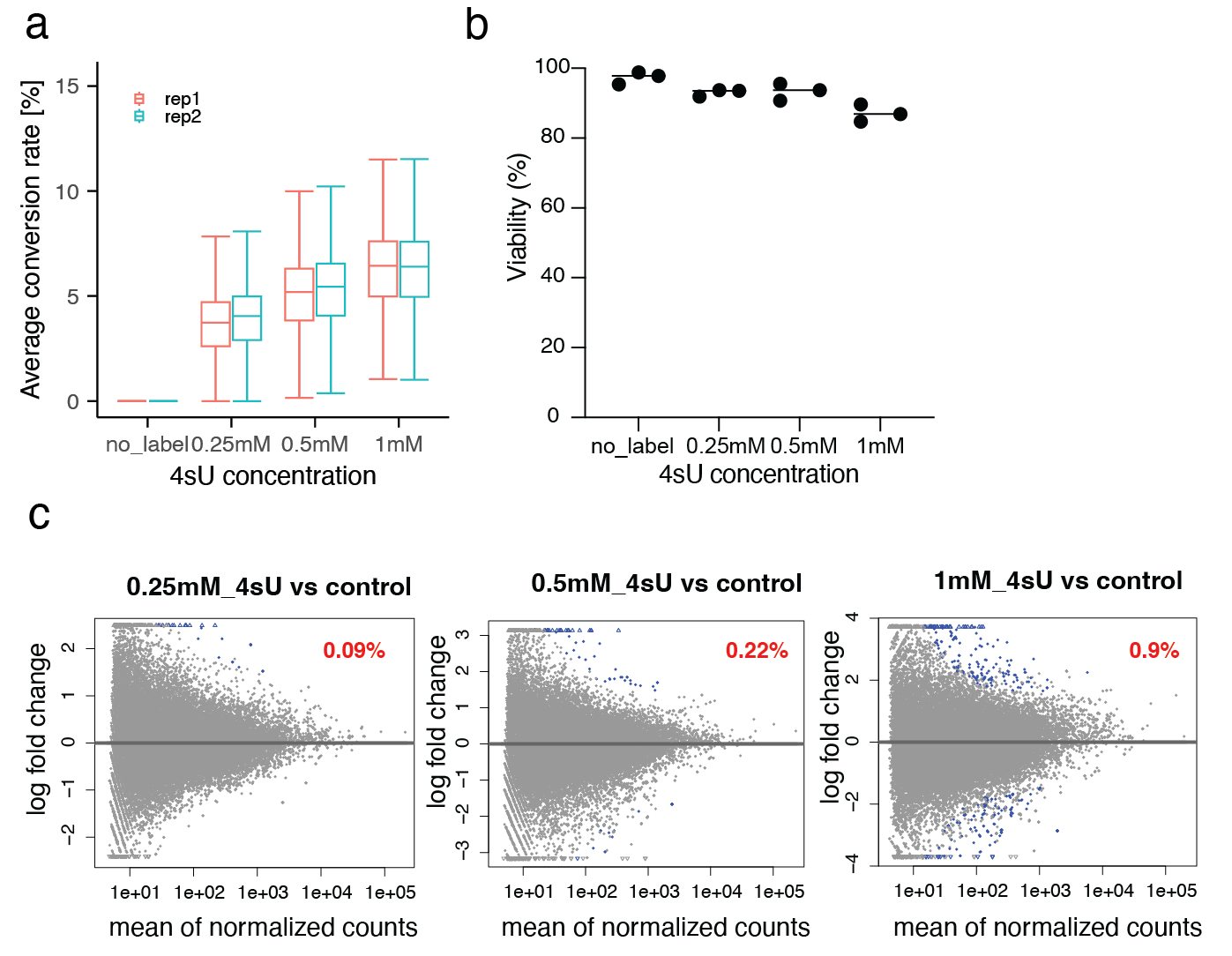

Figure S5. Impact of 4sU labeling on naïve T cells. **a**. T-to-C conversion rates after 4sU labeling using various 4sU concentrations. **b**. Viability of naïve T cells after labeling with various concentrations of 4sU for 1 hour. **c**. Transcriptome changes upon 4sU treatment. The DEGs were colored in blue with their fractions highlighted in red.

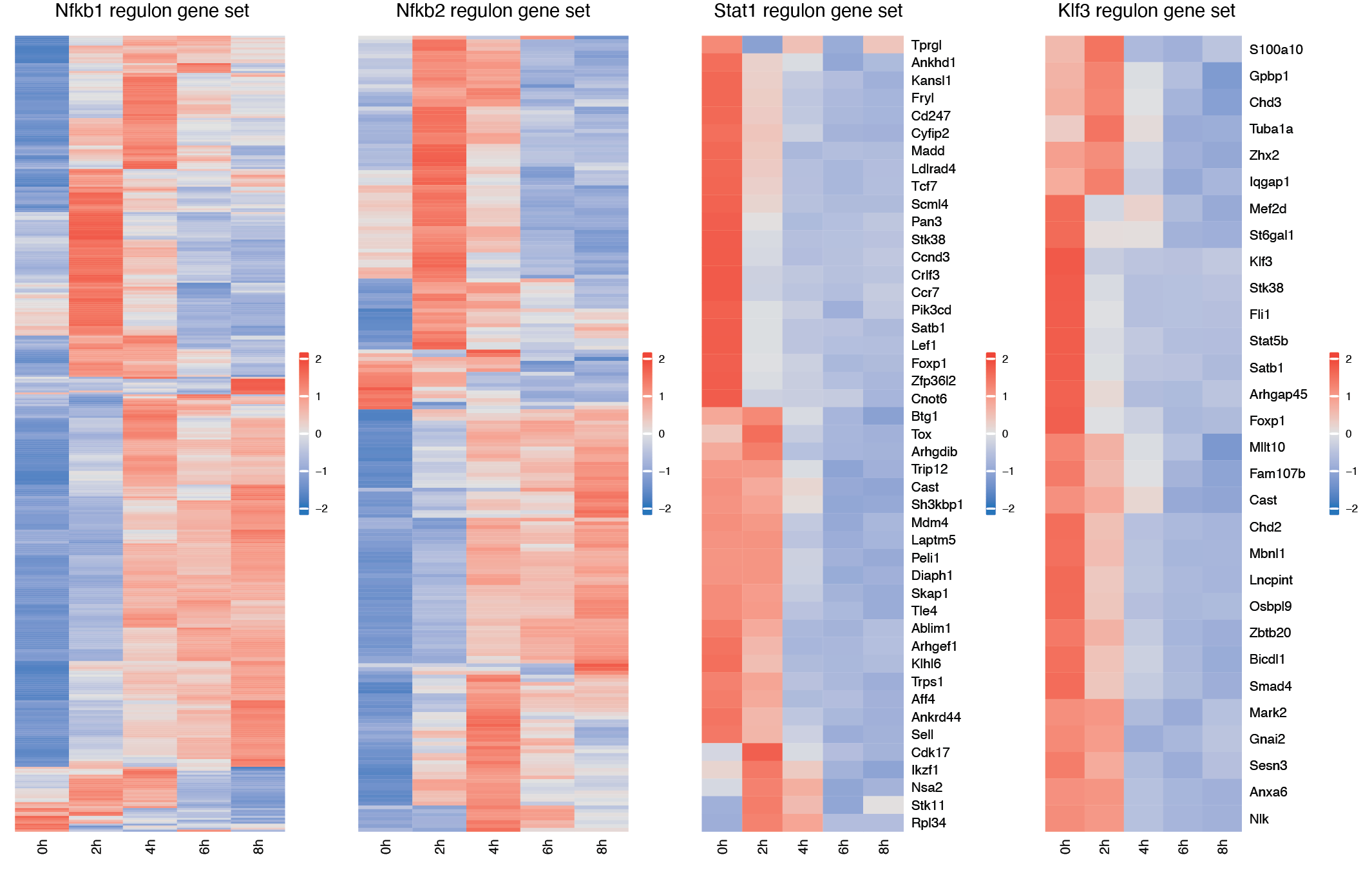

Figure S6. Transcription levels of downstream genes in the regulons of *Nfκb1* (n= 407 genes), *Nfκb2*(n=213 genes), *Stat1*(n= 46 genes) and *Klf3*(n= 30 genes), at different time points during naïve T-cell activation. The data matrix was scaled for each row. Gene names in the Nfkb1 and Nfkb2 regulons were not shown.

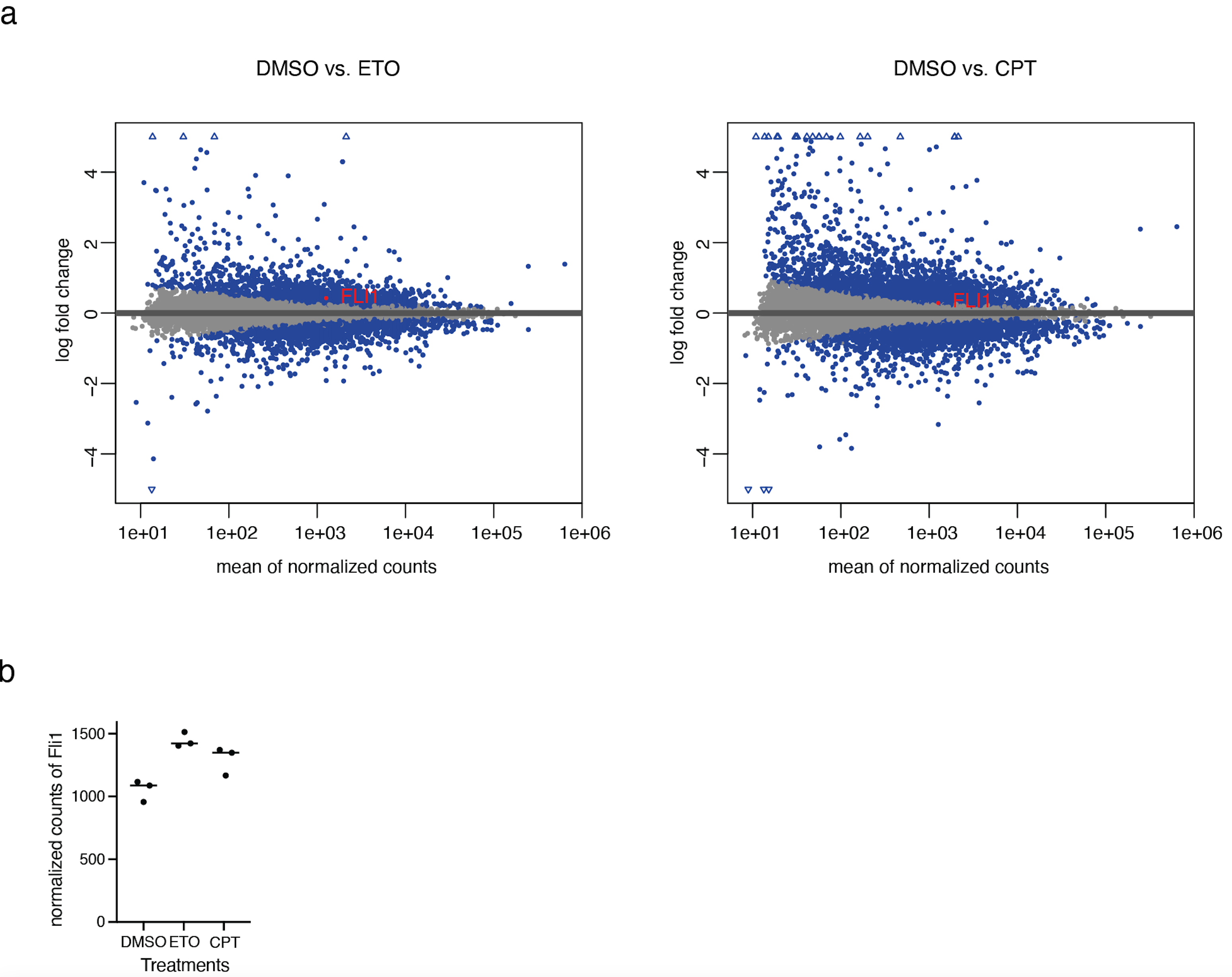

Figure S7. Transcriptome profiling of T cells after topoisomerase inhibition. **a**. Differentially expressed genes after 50 nM CPT (left) or 1 µM ETO (right) treatment for 24h. **b**. Normalized counts of Fli1 total transcripts under indicated treatments.

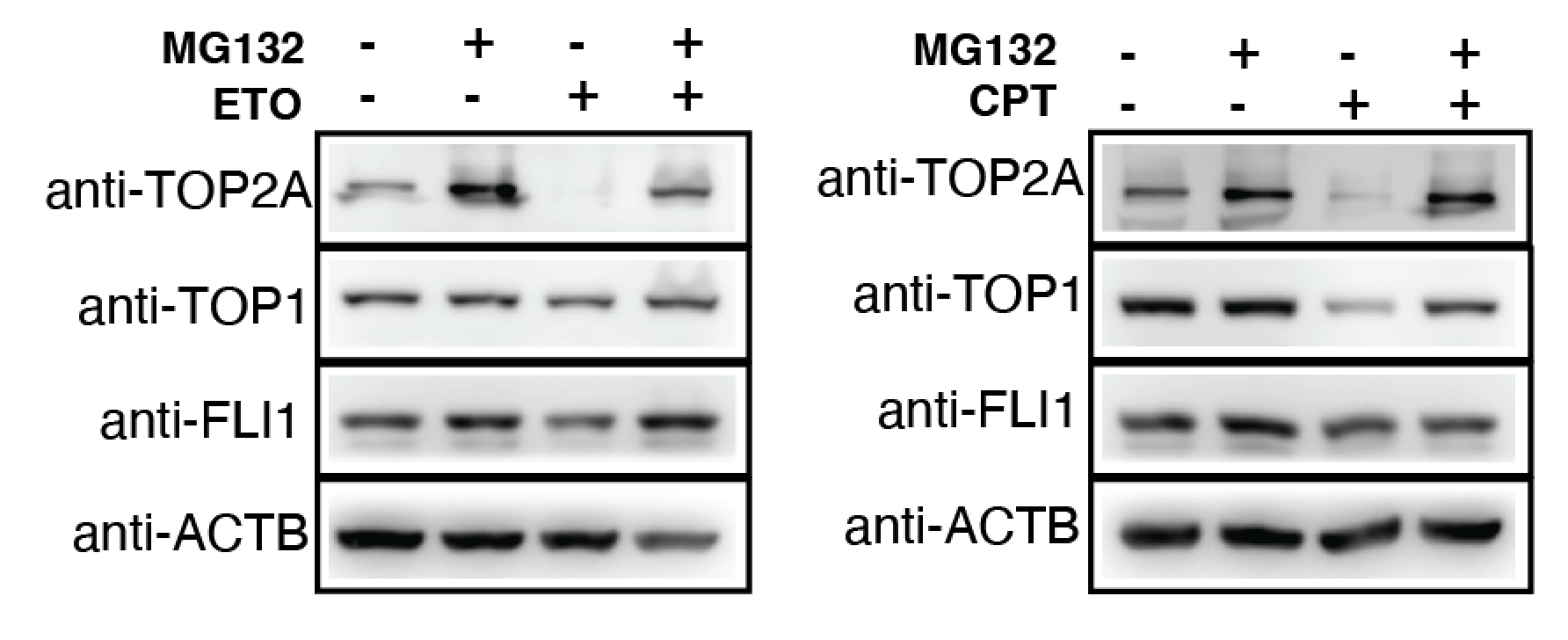

Figure S8. Duplicated Western blot experiment of Fig. 5d to support the proteasome dependence of the degradation of FLI1, TOP1, and TOP2A upon TOPcc formation.
